# An *in vivo* drug repurposing screen and transcriptional analyses reveals the serotonin pathway and GSK3 as major therapeutic targets for NGLY1 deficiency

**DOI:** 10.1101/2021.11.10.468087

**Authors:** Kevin A. Hope, Alexys R. Berman, Randall T. Peterson, Clement Y. Chow

## Abstract

NGLY1 deficiency, a rare disease with no effective treatment, is caused by autosomal recessive, loss-of-function mutations in the *N-glycanase 1 (NGLY1)* gene and is characterized by global developmental delay, hypotonia, alacrima, and seizures. We used an adult *Drosophila* model of NGLY1 deficiency to conduct an *in vivo,* unbiased, small molecule, repurposing screen of FDA-approved drugs to identify therapeutic compounds. Seventeen molecules rescued lethality in a patient-specific NGLY1 deficiency model, including multiple serotonin and dopamine modulators. Exclusive *dNGLY1* expression in serotonin and dopamine neurons, in an otherwise *dNGLY1*-null fly, was sufficient to rescue lethality. Further, genetic modifier and transcriptomic data supports the importance of serotonin signaling in NGLY1 deficiency. Connectivity Map analysis identified glycogen synthase kinase 3 (GSK3) inhibition as a potential therapeutic mechanism for NGLY1 deficiency, which we experimentally validated with TWS119 and lithium. Strikingly, GSK3 inhibitors and a serotonin modulator rescued size defects in *dNGLY1* deficient larvae upon proteasome inhibition, suggesting that these compounds act through NRF1, a transcription factor that regulates proteasome expression. This study reveals the importance of the serotonin pathway in NGLY1 deficiency, and serotonin modulators or GSK3 inhibitors may be effective therapeutics for this rare disease.

## Introduction

NGLY1 deficiency is a rare disease caused by autosomal recessive, loss-of-function mutations in the gene *N-glycanase 1 (NGLY1),* first reported in 2012 (Need et al., 2012). NGLY1 deficiency phenotypes include alacrima, motor impairments, failure to thrive, gastrointestinal issues, developmental delay, hearing loss, and seizures (Enns et al., 2014; Lam et al., 2017). Current treatment options are limited and focus on treating individual symptoms such as lubricating eye drops, feeding therapy, broad-spectrum anti-epileptics, and standard treatments for hearing loss and sleep apnea (Lam et al., 2018). Despite advances in understanding the pathophysiology of NGLY1 deficiency, targeted therapies do not exist for NGLY1 deficient individuals.

The NGLY1 protein is a cytosolic deglycosylase that cleaves N-linked glycans from substrate proteins. Proteins are glycosylated in the endoplasmic reticulum (ER) during folding, and misfolded proteins are retrotranslocated to the cytoplasm (Qi et al., 2017). NGLY1 is believed to play a role in ER-associated degradation (ERAD) based on its physical association with components of the ERAD pathway (Katiyar et al., 2005; McNeill et al., 2004; Park et al., 2001). However, it remains unclear whether NGLY1 plays a critical role in the ER stress response. ER stress markers were elevated in NGLY1 deficient MEFs and RPE-1 cells (Galeone et al., 2020; Lebedeva et al., 2021), yet ER stress was not apparent in other cellular or animal models of NGLY1 deficiency (Asahina et al., 2020; Owings et al., 2018; Tambe et al., 2019). Protein deglycosylation may not be necessary for ERAD, as some NGLY1 substrates are degraded regardless of their glycosylation state (Kario et al., 2008). Alternatively, NGLY1 may deglycosylate cytoplasmic proteins for reasons unrelated to protein degradation.

The first high-confidence NGLY1 substrate identified was the transcription factor NRF1, encoded by the gene *NFE2L1* (Lehrbach & Ruvkun, 2016; Tomlin et al., 2017). NRF1 mediates the proteasome ‘bounce-back’ response, whereby NRF1 upregulates proteasome gene expression upon proteasome inhibition (Radhakrishnan et al., 2010). Under non-stressed conditions, NRF1 is constitutively degraded by the proteasome. NRF1 is deglycosylated by NGLY1 and escapes proteasome degradation during proteasome stress, proteasome inhibition, or increased protein degradation load. NGLY1-dependent deglycosylation activates NRF1, allowing it to enter the nucleus and increase proteasome subunit gene expression (Lehrbach et al., 2019). Inhibition or knockout of NGLY1 increases the level of unprocessed NRF1 and reduces proteasome expression and protein degradation (Tomlin et al., 2017). These data suggest that NGLY1 deficient patients have impaired NRF1 signaling.

The interaction between NGLY1 and NRF1 underlies the observation that NGLY1 deficient cells and animal models display increased sensitivity to proteasome inhibitors (Iyer et al., 2019; Lehrbach & Ruvkun, 2016; Rodriguez et al., 2018; Tomlin et al., 2017), and the regulation of NRF1 by NGLY1 has been the focus of therapeutic development. A screen using *Drosophila* and *C. elegans* that are heterozygous for *NGLY1* mutations and sensitized with the proteasome inhibitor bortezomib found that the atypical antipsychotic aripiprazole restored larval growth defects and upregulated NRF2 in mammalian cells (Iyer et al., 2019), a transcription factor closely related to NRF1. A second potential therapeutic approach has been to target NRF2 more directly. Sulforaphane inhibits KEAP1, an NRF2 inhibitor, subsequently releasing NRF2 from inhibition. NRF2 activation with sulforaphane partially compensated for proteasome transcriptional defects due to NRF1 dysregulation in *Ngly1^-/-^* cells (Yang et al., 2018). While promising, these approaches focused on the known interaction between NGLY1 and NRF1. Previously, our laboratory found that N-acetylglucosamine supplementation rescued lethality in *dNGLY1* deficient *Drosophila* without restoring *cap’n’collar (Drosophila* ortholog of *NRF1*) dependent changes in proteasome gene expression (Owings et al., 2018). Additionally, we recently reported that NGLY1 regulates the glycosylation state and function of NKCC1, a Na-K-2Cl cotransporter (Talsness et al., 2020), but this reduced function was unrelated to the proteasome defects, demonstrating a role for NGLY1 outside of ERAD and proteasome regulation.

NGLY1 is likely involved in a broad range of cellular processes, and exactly how the loss of NGLY1 results in the phenotypes observed in NGLY1 deficiency remains unknown. Previous screens using *Drosophila* focused either on heterozygous animals, larval size, or exposure to proteasome inhibitors in order to focus on the interaction between NGLY1 and NRF1 (Iyer et al., 2019; Rodriguez et al., 2018). Here, we report the first unbiased, phenotypic, drug discovery approach, without focusing on the proteasome, to identify small molecules that rescue lethality in an adult *Drosophila* model of *NGLY1* deficiency. We took a drug repurposing approach and screened the Prestwick Chemical Library, consisting of compounds already approved by the Food and Drug Administration/European Medicines Agency to reduce the time and costs associated with bringing an effective therapeutic to a rare disease population. We identified 17 compounds that rescued lethality in *dNGLY1* deficient flies, four of which modulate serotonin and dopamine signaling. We conducted cell-type-specific rescue experiments and found that exclusive *dNGLY1* expression in serotonin and dopamine neurons was sufficient for rescue in an otherwise *dNGLY1-null* fly. Furthermore, we utilized the Connectivity Map (Lamb et al., 2006; Subramanian et al., 2017), an *in silico,* gene-expression-based screening tool to uncover additional compounds that may be therapeutic for NGLY1 deficiency and identified glycogen synthase kinase 3 (GSK3) as a therapeutic target. Two GSK3 inhibitors, TWS119 and lithium, rescued lethality in *dNGLY1* deficient flies. The GSK3 inhibitors and the serotonin modulator trimipramine rescued proteasome defects observed in *dNGLY1* deficient flies, suggesting that the serotonin and dopamine signaling compounds may act through the GSK3 pathway as well. Our data suggest that GSK3 inhibitors and monoamine signaling modulators, particularly those acting through serotonin and dopamine, may be effective therapeutics for NGLY1 deficiency.

## Results

### Lethality suppression screen identifies 17 primary hit drugs

We developed a small molecule screen to identify compounds capable of rescuing lethality in *dNGLY1^-/-^* flies (Rodriguez et al., 2018) (**Figure 1A**). In our lab, *dNGLY1^-/-^* flies are 100% lethal and fail to emerge from the pupal case. Under normal rearing conditions we have never observed a single adult *dNGLY1^-/-^* fly. To maintain this strain, the null allele is balanced over the *CyO* balancer chromosome that carries a wildtype *dNGLY1* allele, referred to as *dNGLY1^+/-^.* We performed *dNGLY1^+/-^* intercross mass matings and allowed females to lay eggs on drug or DMSO control food. The only flies we expect to observe from these crosses are *dNGLY1^+/-^* because the homozygous nulls *(dNGLY1^-/-^)* and homozygous balanced flies *(Cyo/Cyo)* are lethal. Thus, the appearance of even a single adult *dNGLY1^-/-^* fly was considered a hit in the primary screen. We screened 1,040 compounds from the Prestwick Chemical Library, consisting of 20,108 flies, and identified 17 hit compounds, a hit rate of 1.6% (**Figure 1B, Table S1**). In all DMSO treated controls, we observed 1,658 *dNGLY1^+/-^* and zero *dNGLY1^-/-^* adult flies. Hit compounds fell into five broad categories. The first category was iodine-containing contrast imaging agents, consisting of iocetamic acid, iodixanol, and ioversol. The second class of compounds were antibiotics and comprised of moxalactam and pththalylsulfathiazole. The third class included the three anti-inflammatory agents mesalamine, dichlorphenamide, and nimesulide. The fourth class contained compounds that modulate monoamine signaling, particularly the dopaminergic and serotoninergic neurotransmitter systems, including urapidil, trimipramine, ipsapirone, and bromocriptine. Five compounds did not fall into a discrete group and included nifekalant, two stereoisomers methylhydantoin-5-(L) and methylhydantoin-5-(D), theobromine, and nocodazole.

**Figure 1.**
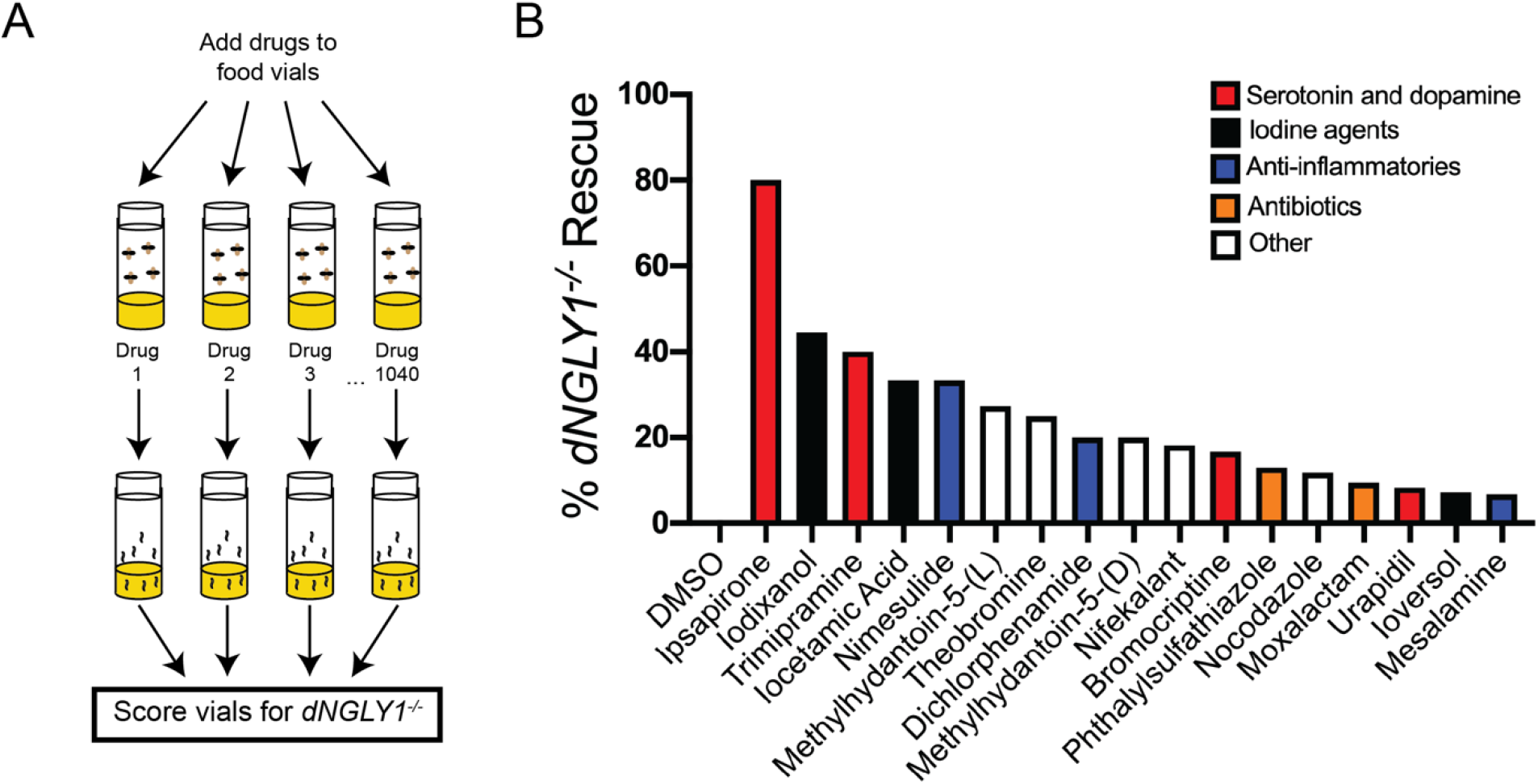
Workflow and results of the *in vivo,* small molecule screen. **A)** Screening strategy to identify compounds that rescue lethality in *dNGLY1^-/-^* flies. **B)** Seventeen hit compounds were identified from the 1040 compounds screened from the Prestwick Chemical Library. Percent expected *dNGLY1^-/-^* was calculated from each drug treatment based on the number of heterozygous *dNGLY1^+/-^* flies that emerged from each vial.

Seven compounds resulted in no adults of any genotype (**Table S1**), potentially indicating lethality to heterozygous *dNGLY1^+/-^* flies or flies in general. Avermectin B1 and ivermectin are pesticides and likely kill all *Drosophila,* regardless of *dNGLY1* genotype. Additional compounds resulting in lethality in *dNGLY1^+/-^* were the dihydrofolate reductase inhibitor methotrexate, the nicotinic acetylcholine receptor modulator galanthamine hydrobromide, the non-selective monoamine oxidase inhibitor nialamide, the β_2_ adrenergic receptor agonist levalbuterol, and the anticholinergic compound tridihexethyl chloride. While we did not further explore these compounds in this study, they may reveal new aspects of NGLY1 biology.

### Transcriptomic and genetic data implicate the serotonin and dopamine system in NGLY1 deficiency

To prioritize candidate small molecules for follow-up experiments, we reanalyzed RNAseq data previously published by our laboratory in which we characterized *dNGLY1* RNAi knockdown flies (Owings et al., 2018). We observed a significant downregulation of nearly the entire dopaminergic biosynthesis pathway (**Figure 2A**), including the enzyme *Tyrosine Hydroxylase,* or *Pale* in the fly (log2FoldChange = −1.40, padj = <0.00001), the dopamine transporter *DAT* (log2FoldChange = −0.65, padj = 0.0033), *Ebony* (log2FoldChange = −0.39, p = 0.018), and upregulation of *yellow* (log2FoldChange = 2.15, padj = <0.00001). We also observed the downregulation of a host of enzymes in the tyrosine breakdown pathway that converts tyrosine to fumarate and acetoacetate (**Figure 2A**).

**Figure 2.**
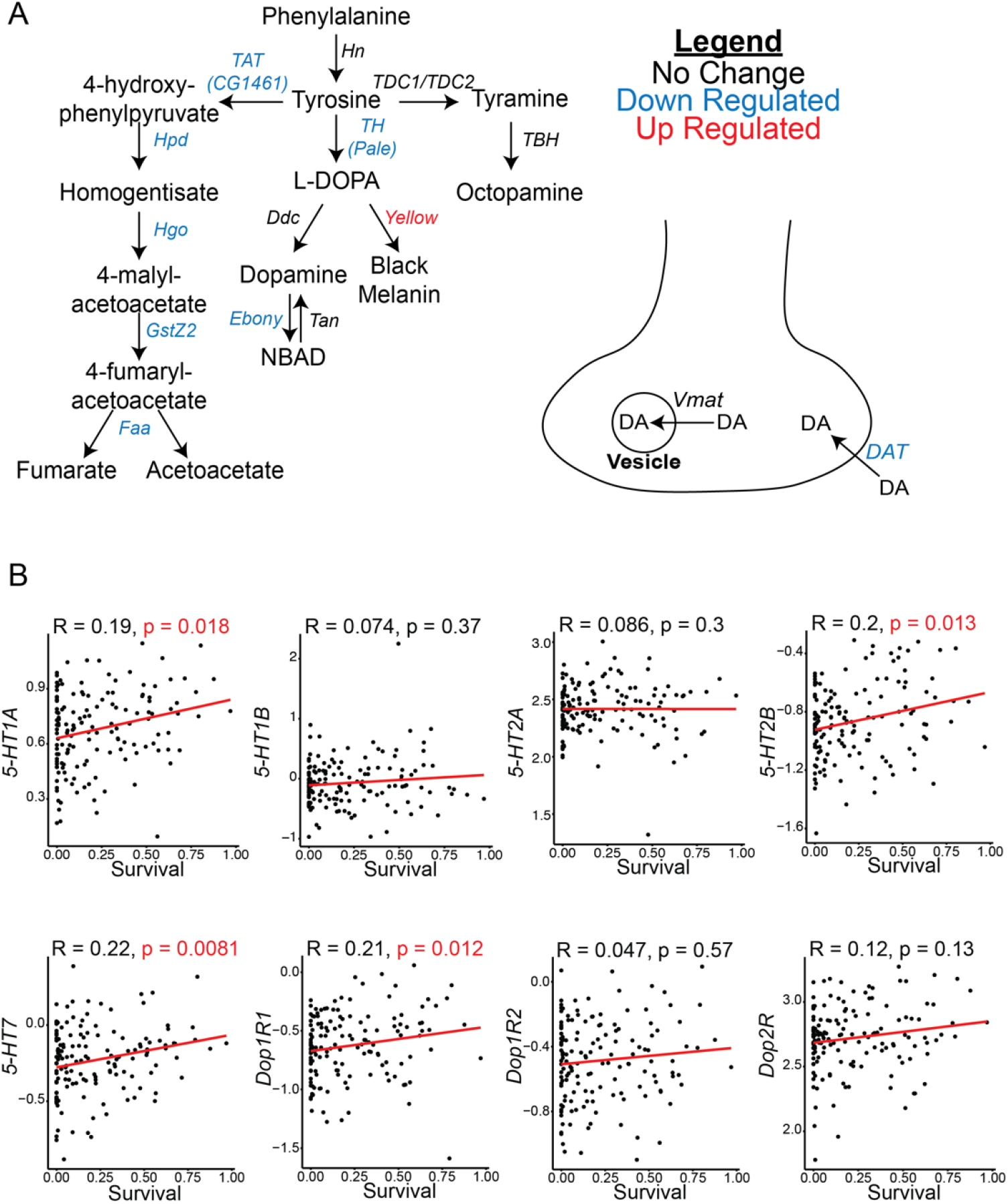
Prior studies implicate the serotonin and dopamine system in NGLY1 deficiency. **A)** Diagram showing the differentially expressed genes in the *Drosophila* dopamine biosynthesis and transport pathways. Gene expression data was from *Tubulin>dNGLY1-RNAi* whole flies reported in a previous study (Owings et al., 2018). **B)** Spearman correlation analysis between survival upon knockdown of *dNGLY1* (Talsness et al., 2020) and gene expression of serotonin and dopamine receptors at baseline in the Drosophila Genetic Reference Panel (Everett et al., 2020).

Our lab previously performed a genetic modifier screen for genes that impacted lethality caused by *dNGLY1* RNAi knockdown in the Drosophila Genetic Reference Panel (DGRP) (Talsness et al., 2020). We identified 61 genes that were associated with survival upon knockdown of *dNGLY1,* and one of the top hits was the serotonin receptor *5-HT1A.* We next examined whether expression levels of *5-HT1A* correlated with survival upon knockdown of *dNGLY1.* Using a previously published DGRP RNAseq dataset (Everett et al., 2020), we performed a correlation analysis between *5-HT1A* expression levels at baseline in the DGRP and survival for each DGRP strain upon knockdown of *dNGLY1* (**Figure 2B**). We observed a positive correlation between *5-HT1A* expression levels and survival (Rho = 0.193, p = 0.01768, Spearman Rank Correlation). We also examined correlations between survival upon *dNGLY1* knockdown in the DGRP and expression levels for all *Drosophila* serotonin and dopamine receptors. Significant positive correlations were observed between the *5-HT2B* (Rho = 0.2, p = 0.013), *5-HT7* (Rho = 0.22, p = 0.0081), and *Dop1R1* (Rho = 0.21, p = 0.012) receptors (**Figure 2B)**. Together, these transcriptomic and genetic data provide further support that the serotonin and dopaminergic pathways are important to NGLY1 deficiency and may explain why molecules targeting these pathways rescue NGLY1 deficiency.

### Secondary confirmation of serotonin, dopamine, and anti-inflammatory compounds identified in the primary screen

We prioritized the serotonin and dopamine signaling compounds based on the previous data described above implicating the serotonin and dopamine system. We retested ipsapirone, bromocriptine, urapidil, and trimipramine at the same concentration as the primary screen and concentrations five times higher and lower in triplicate. We tested urapidil at only 2.5 times higher due to lower solubility levels. We observed a dose-dependent rescue effect for all four compounds. Ipsapirone showed 2.6% rescue at 1 μM (n = 1/38; observed/expected; **Figure 3A, Table S2**), 19.5% at 5 μM (n = 8/41), and 21.5% at 25 μM (n = 7/33). Trimipramine displayed 0% (n = 0/55) rescue at 1 μM, 7.8% (n = 3/39 at 5 μM, and 17.6% (n = 6/34) at 25 μM. Urapidil rescued lethality to 11.5% (n = 5/44) at 1 μM, 8.9% (n = 4/45) at 5 μM, and to 4.5% (n = 2/44) at 12.5 μM. Finally, for bromocriptine we observed a 4.1% (n = 2/49) rescue at 1 μM, 7.0% (n = 3/43) rescue at 5 μM, and 0% (n = 0/45) rescue at 25 μM. These data confirm these serotonin and dopamine signaling compounds as hits from our primary screen.

**Figure 3.**
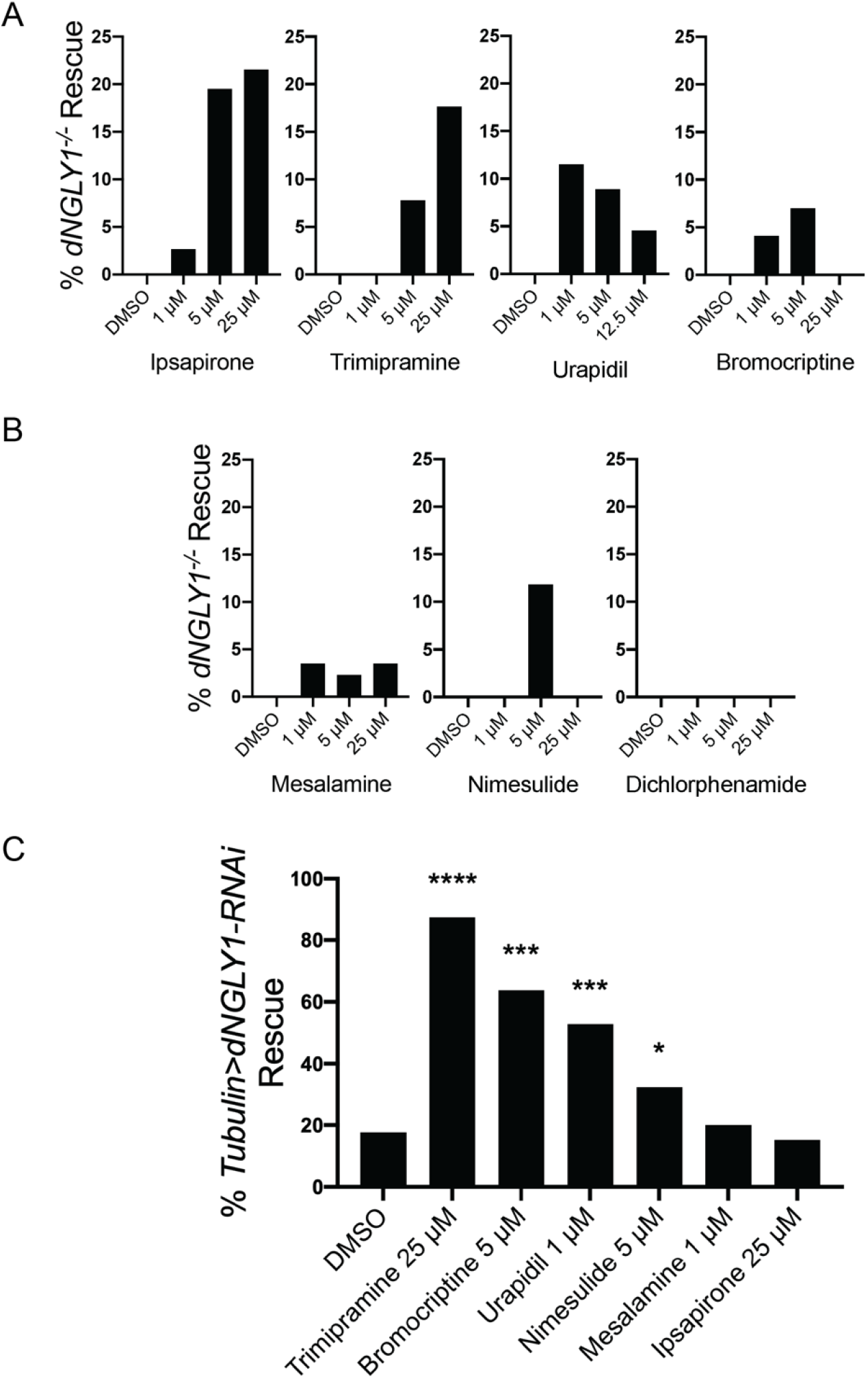
Secondary confirmation of serotonin, dopamine, and anti-inflammatory compounds. Each compound was retested in a secondary confirmation assay at 1 μM, 5 μM, and 25 μM. **A)** Ipsapirone, trimipramine, urapidil, and bromocriptine have mechanisms of action through the serotonin and/or dopamine signaling pathways and all showed rescue effects in the secondary confirmation. **B)** Mesalamine, nimesulide, and dichlorphenamide are classified as anti-inflammatory compounds. While mesalamine and nimesulide showed rescue, dichlorphenamide failed to show rescue in a secondary confirmation. **C)** Drugs that rescued lethality in the secondary assay were tested at the most effective concentration using a separate NGLY1 deficiency model, *Tubulin>dNGLY1-RNAi.* On DMSO, 17.7% of the expected number of *Tubulin>dNGLY1-RNAi* flies emerge. While trimiprimine, bromocriptine, urapidil, and nimesulide showed rescue, mesalamine, and ipsapirone failed to rescue. **** p < 0.0001, *** p < 0.001, and * p < 0.05.

Based on known safety profiles and the potential for use in a pediatric population, we also followed up on the anti-inflammatory compounds. Two of the three compounds confirmed rescue of lethality in *dNGLY1^-/-^* flies in secondary validation assays. Mesalamine rescued lethality to 3.5% (n = 1/76.5; **Figure 3B**), 2.3% (n = 1/44), and 3.5% (n = 1/29) at 1 μM, 5 μM, and 25 μM, respectively. Nimesulide only showed rescue at 5 μM (11.8%; n = 3/26), with no rescue at 1 μM (n = 0/48) or 25 μM (n = 0/72). Dichlorphenamide did not rescue lethality at any concentration (1 μM: n = 0/71; 5 μM: n = 0/60; 25 μM: n = 0/36).

To determine whether hit compounds from our screen rescued lethality in another *Drosophila* model of *NGLY1* deficiency, we tested the compounds in the *Tubulin>dNGLY1-RNAi* model previously characterized by our lab (Owings et al., 2018) (**Figure 3C, Table S2**). Unlike the *dNGLY1^-/-^* flies, *Tubulin>dNGLY1-RNAi* flies are not completely lethal and 17.7% of the expected *Tubulin>dNGLY1-RNAi* flies survive to adulthood on DMSO control food (n = 23/130; **Figure 3C, Table S2**). We tested compounds at the concentrations with the highest rescue in the *dNGLY1-*null model. Three out of four of the serotonin and dopamine hits rescued lethality in *Tubulin>dNGLY1-RNAi* flies, compared to DMSO control. Trimipramine rescued *Tubulin>dNGLY1-RNAi* flies to 87.5% (n = 35/40, x^2^ = 26.55, p < 0.00001), bromocriptine rescued to 63.8% (n = 37/58, x^2^ = 18.28, p < 0.001), and urapidil rescued to 52.8% of the expected number (n = 28/53, x^2^ = 11.86, p < 0.001) (**Figure 3C, Table S2**). Ipsapirone did not rescue in *Tubulin>dNGLY1-RNAi* at 25 μM (n = 16/105, x^2^ = 0.181, p = 0.67). The antiinflammatory compound nimesulide at 5 μM (n = 24/74, x^2^ = 4.977, p = 0.026) rescued, but mesalamine at 1 μM (n = 15/75, x^2^= 0.115, p = 0.735) did not rescue lethality in the *Tubulin>dNGLY1-RNAi* model (**Figure 3C, Table S2**). Testing hit drugs from our primary and secondary screen in a different *Drosophila* model provides further evidence that the small molecules acting through the serotonin and dopamine signaling pathways are beneficial to NGLY1 deficiency.

### *dNGLY1* expression in serotonin and dopamine neurons rescues lethality

Since we observed multiple hit drugs in the serotonin and dopamine signaling pathways and transcriptomic and genetic data implicating these pathways, we determined which serotonergic and dopaminergic cell types require *dNGLY1* expression for survival. We sought to express wildtype *dNGLY1* in serotonergic and dopaminergic neurons in an otherwise *dNGLY1* null fly. First, we tested whether expression of *dNGLY1* in the same pattern as endogenous *dNGLY1* could rescue lethality. We utilized a *dNGLY1^ΔGAL4^* line that harbors a cassette inserted into the endogenous *dNGLY1* locus carrying a stop codon followed by the GAL4 coding sequence to generate a *dNGLY1*-null allele while expressing *GAL4* in the endogenous *dNGLY1* pattern (P.-T. Lee et al., 2018). Thus, *dNGLY1^ΔGAL4^* can express any UAS-transgene in the same pattern as endogenous *dNGLY1.* We generated *dNGLY1*-null flies *(dNGLY1^ΔGAL4/-^*) that expressed wildtype *UAS-dNGLY1^wt^* (Funakoshi et al., 2010) in the endogenous pattern. We observed full rescue of *dNGLY1^ΔGAL4^* flies carrying the *UAS-dNGLY1^wt^* transgene (**Figure 4, Table S3**). However, as expected, *UAS-dNGLY1^C303A^* (Funakoshi et al., 2010), a catalytically inactive form of *dNGLY1,* failed to rescue lethality. These data indicate that expressing wildtype *dNGLY1* with the GAL4/UAS system can rescue lethality in *dNGLY1* null flies.

**Figure 4.**
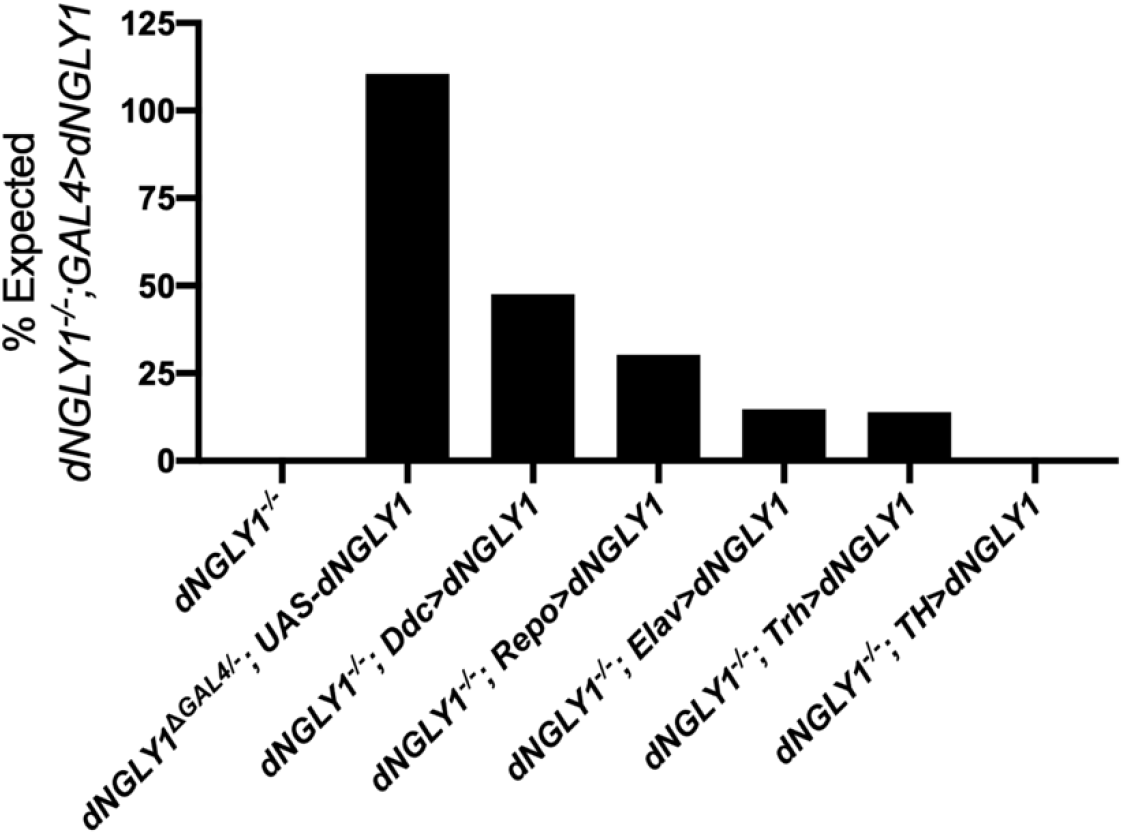
Expressing *dNGLY1* in dopamine and/or serotonin neurons rescues lethality in *dNGLY1^-/-^* flies. The GAL4/UAS system was used to express *dNGLY1* in a cell-type specific pattern in otherwise *dNGLY1^-/-^* flies. Expressing *dNGLY1* in the endogenous expression pattern with *dNGLY1^ΔGAL4^* completely rescued the lethality phenotype of *dNGLY1^-/-^* flies. Expressing *dNGLY1* only in dopamine and serotonin neurons with *Ddc-GAL4* rescued to 47.5%. *dNGLY1* expression in glial cells with *Repo-GAL4* rescued lethality to 30.3%, while expression in all neurons with *Elav-GAL4* rescued to 14.7%. Expressing *dNGLY1* in only serotonin neurons with *Trh-GAL4* rescued to 13.8%, and expression in only dopamine neurons with *TH-GAL4* did not rescue lethality.

Next, we tested whether serotonin- and dopamine-related cell types require *dNGLY1* expression for survival using cell-type-specific GAL4 lines. To do this, we generated *dNGLY1^-/-^* flies (same strain used in drug screen with 100% lethality) that carried different cell-specific GAL4 transgenes and the *UAS-dNGLY1* transgenes tested above. Expression of *dNGLY1^wt^* in all neurons *(Elav-GAL4)* or all glia *(Repo-GAL4)* resulted in 14.7% (n = 13/89) and 30.3% (n = 18/60) rescue of lethality, respectively. Expression of *dNGLY1^wt^* in neurons or glia is sufficient for partial rescue of lethality in an otherwise *dNGLY1* null fly. Expression of *dNGLY1^wt^* simultaneously in both dopamine and serotonin neurons *(Ddc-GAL4)* rescued lethality to 47.5% (n = 29/61). *dNGLY1^wt^* expression in dopamine neurons only *(TH-GAL4)* failed to rescue (n = 0/37). However, *dNGLY1^wt^* expression in serotonin neurons *(Trh-GAL4)* rescued lethality to 13.8% (n = 10/72). It is possible that some variation in rescue efficacy may be due to differences in the strength of GAL4-induced transgene expression. Expressing the catalytically dead *dNGLY1^C303A^* did not rescue lethality in any cell type (**Table S3**), suggesting that the catalytic activity of dNGLY1 is necessary for the rescue effect. These data indicate that *dNGLY1* activity in both dopamine and serotonin neurons is sufficient for survival in an otherwise *dNGLY1-null* fly, and serotonin neurons may be the predominant cell type underlying the rescue effect.

### Connectivity Map identifies GSK3 inhibition as a therapeutic mechanism

In addition to our *in vivo* small molecule screen, we used the Connectivity Map (CMAP), an *in silico,* gene expression-based tool to identify potential therapeutic compounds (Lamb et al., 2006; Subramanian et al., 2017). We used our previously published RNAseq dataset on *Tubulin>dNGLY1-RNAi* flies (Owings et al., 2018) to identify compounds that reverse the transcriptional effects caused by reduced levels of dNGLY1. We selected the top 200 upregulated and 200 downregulated genes from *Tubulin>dNGLY1-RNAi* flies, identified the human orthologs, and filtered for conserved genes between flies and humans (DIOPT score > 7). These criteria yielded 95 upregulated and 87 downregulated genes to input into the CMAP pipeline (**Table S4**). Using an enrichment score cutoff of >90 or <-90, we identified 10 compounds predicted to cause transcriptional effects similar to *dNGLY1* knockdown and 30 compounds predicted to cause transcriptional effects that counteract the loss of *dNGLY1* (**Figure 5A, B**). There was an enrichment in the 10 compounds that mimic the effect of *dNGLY1* knockdown in the pharmacologic class “Proteasome Inhibitor” (Score = 94.47; **Table S4**). Additionally, we capitalized on a gene expression data set generated from a liver-specific knockout of *Ngly1* in the mouse (Fujihira et al., 2020). CMAP analysis of differentially expressed genes from *Ngly1* liver-specific knockout revealed 193 compounds predicted to cause similar transcriptional effects as the loss of *Ngly1* and 195 compounds predicted to counteract *Ngly1* loss. Compounds predicted to mimic *Ngly1* loss were similarly enriched in “Proteasome Inhibitor” (Score = 99.33) (**Table S4**). Among the compounds predicted to mimic the effects of reduced *dNGLY1/Ngly1,* six were shared between the fly and liver datasets (**Figure 5A**). Three of the shared mimic compounds, MLN-2238, MG-132, and NSC632839 inhibit the ubiquitin-proteasome system. This is not surprising as the proteasome is downregulated in both models because of the NGLY1-dependent regulation of NRF1. The glycogen synthase kinase inhibitor TWS119 was the only overlapping compound predicted to counteract the transcriptional effect in both data sets (**Figure 5B**).

**Figure 5.**
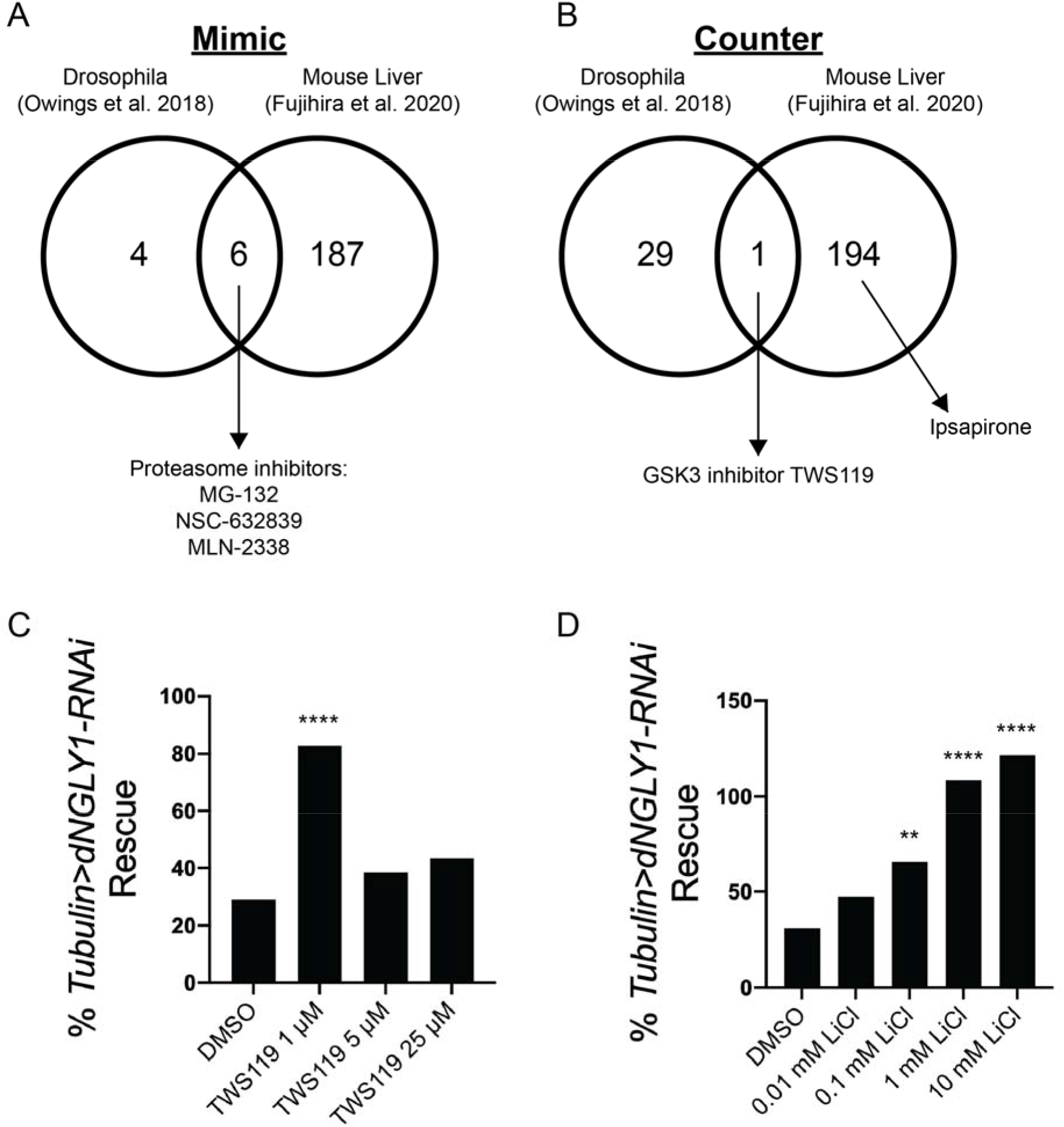
CMAP analyses identifies GSK3 inhibition as a therapeutic mechanism. **A)** Ten compounds were predicted to mimic the transcriptional response due to *dNGLY1* knockdown in flies (Owings et al., 2018). In the mouse liver data set, 193 compounds were predicted to mimic the transcriptional response due to the loss of *Ngly1* (Fujihira et al., 2020). The fly and mouse data sets shared six compounds, three of which are proteasome inhibitors. **B)** In the fly dataset, 30 compounds were predicted to counter the transcriptional response due to *dNGLY1* knockdown. In the mouse liver dataset, 195 compounds were predicted to counter the transcriptional response due to loss of *Ngly1.* The GSK3 inhibitor TWS119 was the only compound shared between the two data sets. **C)** TWS119 rescued the lethality of *Tubulin>dNGLY1-RNAi* flies at 1 μM but did not rescue lethality at 5 or 25 μM. **D)** Lithium chloride (LiCl) showed a dose response effect and significantly rescued lethality in *Tubulin>dNGLY1-RNAi* flies at 0.1 mM, 1 mM, and 10 mM. The lowest dose at 0.01 mM did not increase survival of *Tubulin>dNGLY1-RNAi* flies. **** p < 0.0001, ** p < 0.01.

Drugs, like TWS119, predicted to counter gene expression changes associated with NGLY1 deficiency may have therapeutic potential. We experimentally tested whether TWS119 rescued lethality in *Tubulin>dNGLY1-RNAi* flies. We observed a significant increase in the proportion of *Tubulin>dNGLY1-RNAi* flies that eclosed on food containing 1 μM TWS119 compared to DMSO alone (**Figure 5C**, **Table S5**, x^2^ = 15.72, p < 0.001), while higher doses did not show improved survival. In addition to TWS119, we tested whether lithium, the only FDA-approved GSK3 inhibitor, could similarly rescue lethality in *Tubulin>dNGLY1-RNAi* flies. Lithium chloride (LiCl) significantly rescued lethality in *Tubulin>dNGLY1-RNAi* flies at 0.1 mM (x^2^ = 7.86, p = 0.0051), 1 mM (x^2^ = 23.48, p <0.00001), and 10 mM (x^2^ = 29.33, p < 0.00001) (**Figure 5D**, **Table S5**). Low dose LiCl at 0.01 mM had no effect on survival in *Tubulin>dNGLY1-RNAi* flies (x^2^ = 2.30, p = 0.13). We tested lithium acetate to determine if a different source of lithium similarly rescued lethality in *Tubulin>dNGLY1-RNAi* flies and observed similar results as with LiCl. Lithium acetate rescued lethality a 0.1 mM (**x**^2^ = 10.04, p = 0.002) and at 1 mM (x^2^ = 7.238, p = 0.0071) (**Figure S1**, **Table S5**). These data suggest that inhibiting GSK3 may be an effective therapeutic strategy for NGLY1 deficiency.

### GSK3 inhibitors and trimipramine rescue size defects in *dNGLY1^+/-^* larvae upon proteasome inhibition

Previous studies demonstrated that heterozygous *dNGLY1^+/-^* larvae have increased sensitivity to proteasome inhibitors and decreased larval size upon proteasome inhibition (Iyer et al., 2019). To determine whether the GSK3 inhibitors prevent increased sensitivity to proteasome inhibition, we tested whether GSK3 inhibitors could overcome larval size defects observed in *dNGLY1^+/-^* larvae treated with the proteasome inhibitor bortezomib (BTZ). We also tested the serotonin modulator trimipramine since it was a confirmed hit from the primary screen and displayed the strongest rescue in the *Tubulin>dNGLY1-RNAi* model. Confirming the previous study (Iyer et al., 2019), *dNGLY1^+/-^* larva treated with 5 μM BTZ at the 3^rd^ instar stage were significantly smaller than DMSO treated controls (One-way ANOVA, p < 0.0001, Dunnett’s multiple comparison test, p < 0.0001) (**Figure 6A, B**). To test the effect of our compounds, we raised *dNGLY1^+/-^* larvae on food supplemented with each compound, similar to what was done in the drug screen. At the 3^rd^ instar stage, larvae were transferred to food that contained each compound and BTZ. Pretreatment with 1 μM TWS119 (p = 0.24), 1 mM LiCl (p = 0.76), or 25 μM trimipramine (p = 0.58) all rescued larval size defects caused by inhibition of the proteasome with BTZ and were not significantly different from *dNGLY1^+/-^* larvae on DMSO control food. For the most part, these treatments had no effect on wildtype larvae with the exception of BTZ and LiCl pretreatment, where wildtype larvae showed a slight decrease in size (One-way ANOVA, F = 10.42, p < 0.0001) (**Figure 6A and S2**), opposite of what we observed in *dNGLY1^+/-^* larvae. These data suggest that GSK3 inhibitors and trimipramine rescue proteasome defects associated with loss of NGLY1 function.

**Figure 6.**
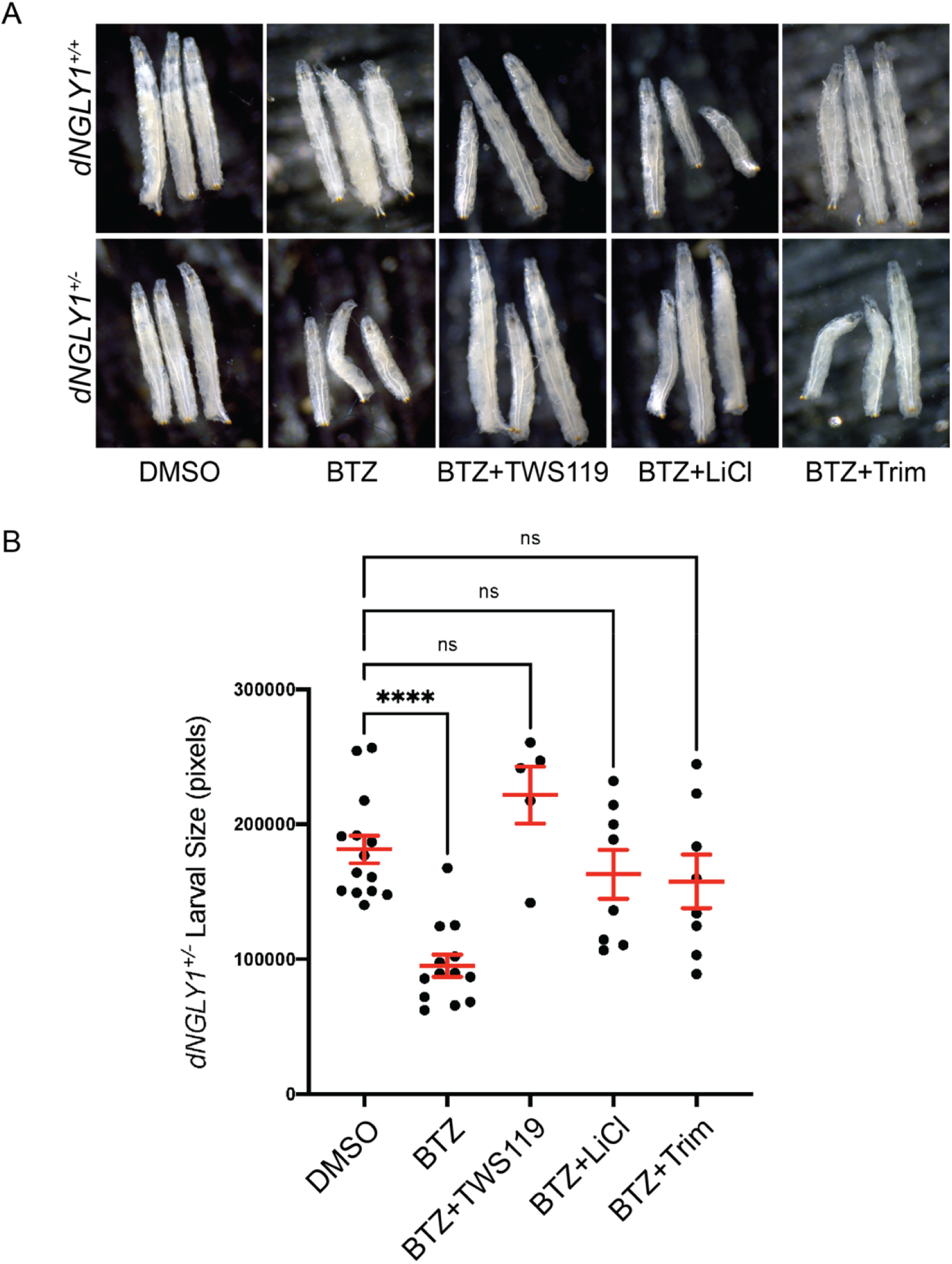
GSK3 inhibitors and trimipramine rescue size defects in *dNGLY1^+/-^* larvae upon proteasome inhibition. Pretreatment with TWS119, LiCl, or trimipramine rescue proteasome inhibition-induced size reduction in *dNGLY1^+/-^* larvae. **A)** Images of *dNGLY1^+/+^* and *dNGLY1^+/-^* treated with DMSO, bortezomib (BTZ) alone, or BTZ with drug. **B)** Quantification of *dNGLY1^+/-^* larval size. *dNGLY1^+/-^* were significantly smaller when treated with 5 μM BTZ compared to DMSO. There were no significant differences in dNGLY1^+/-^ larvae treated with DMSO compared to 5 μM BTZ+1 μM TWS119, 5 μM BTZ+1 mM LiCl, or 5 μM BTZ+25 μM trimipramine, indicating a strong rescue of the susceptibility to proteasome inhibition. BTZ = bortezomib, Trim = trimipramine. Dots represent individual larva and red bars are mean ± SEM. ****p < 0.0001.

## Discussion

Rare diseases, like NGLY1 deficiency, often lack effective therapies. Since the initial discovery of NGLY1 deficiency in 2012, the patient population has grown, along with the body of research. The understanding of the biological function of NGLY1, and the consequences resulting from loss of NGLY1 activity, has expanded in recent years. However, despite this improved understanding, targeted therapeutics for NGLY1 deficiency remain elusive. Here, we utilized *Drosophila* to combine the benefits of phenotypic drug discovery with drug repurposing to identify small molecule therapeutics for NGLY1 deficiency.

This small molecule screen identified four serotonin and dopamine signaling compounds that rescued lethality in *dNGLY1*-null flies, indicating that these molecules bypass the need for *dNGLY1* function for survival. Urapidil is an agonist of the 5-HT_1A_ receptor and an antagonist of the α1-adrenoceptor (Buch, 2010). Ipsapirone is a 5HT_1A_ partial agonist (Lesch et al., 1990; Newman-Tancredi et al., 1998), while trimipramine displays modest inhibition of the serotonin reuptake transporter (Haenisch et al., 2011), as well as antagonistic activity at the 5HT_2A_ receptor, among others (Gross et al., 1991). Additionally, we identified bromocriptine which targets a wide range of receptors, acting as an agonist at multiple dopamine and serotonin receptors, including the dopamine D_2_ receptor and 5HT_1A_, as well as an α-adrenergic receptor antagonist. (Newman-Tancredi, Cussac, Audinot, et al., 2002; Newman-Tancredi, Cussac, Quentric, et al., 2002). Previous work found that aripiprazole, a compound with a pharmacological profile similar to bromocriptine, rescued larval size defects upon inhibition of the proteasome in NGLY1^+/-^ flies and worms (Iyer et al., 2019). Based on the combined pharmacology of these drugs, along with previous work from our lab that found that SNPs in the 5HT_1A_ receptor were associated with survival upon knockdown of *dNGLY1* (Owings et al., 2018), the 5HT_1A_ receptor may be a critical therapeutic target in NGLY1 deficiency. In addition to identifying small molecules that act through serotonin/dopamine signaling, we found that expressing *dNGLY1* in serotonin and dopamine neurons or exclusively in serotonin-producing neurons rescued lethality in an otherwise *dNGLY1-*null fly. These results suggest that NGLY1 activity in the monoamine system is critical for proper development, and the loss of NGLY1 function in serotonergic and dopaminergic neurons drives a significant component of the lethality observed in *dNGLY1* deficient flies. Further studies are needed to determine the receptor, or combination of receptors, that underlie the mechanism of action for these monoamine signaling compounds.

Similar to our findings in *Drosophila,* impairments in the serotonergic and dopaminergic systems likely underlie a large portion of NGLY1 deficiency phenotypes in humans. NGLY1 deficient individuals display tremors, hyperkinesis, and choreiform movements. Cerebrospinal fluid analysis revealed reduced levels of the dopamine and serotonin synthesis cofactor tetrahydrobiopterin, the dopamine metabolite homovanillic acid, and the serotonin metabolite 5-hydroxyindolacetic acid (Lam et al., 2017). An additional case study identified low levels of the serotonin precursor 5-hydroxytryptophan (Lipari Pinto et al., 2020). These studies suggest that dopamine and serotonin signaling is disrupted in NGLY1 deficient individuals. Small molecules that act through serotonin and/or dopamine signaling may be effective therapeutic avenues for NGLY1 deficiency.

Using a complementary approach to our *in vivo,* small molecule repurposing screen, we used the CMAP (Lamb et al., 2006; Subramanian et al., 2017), an *in silico,* gene-expression-based tool to identify compounds that reverse transcriptional defects upon the loss of *NGLY1.* Inhibiting GSK3 using either TWS119 or lithium rescued lethality in *dNGLY1* knockdown flies and larval size defects induced by proteasome inhibition. Previously, our lab reported downregulation of genes regulated by the cap’n’collar (cnc) transcription factor (NRF1/NRF2 in mammals) in flies with reduced levels of *dNGLY1* (Owings et al., 2018). Lithium induces the expression of cnc response genes in the fly through the inhibition of *shaggy,* the *Drosophila* ortholog of GSK3 (Castillo-Quan et al., 2016). NGLY1 regulates NRF1 activity through the cleavage of N-glycans, allowing NRF1 to become active and enter the nucleus (Lehrbach et al., 2019; Tomlin et al., 2017). GSK3 regulates NRF1/NRF2 through an alternative, phosphorylation-dependent mechanism, and targeting GSK3 may bypass the dysregulation of NRF1 due to the absence of NGLY1. In *C. elegans,* the NRF1/NRF2 ortholog *SKN-1* is phosphorylated and inhibited by GSK-3 (An et al., 2005). In mammalian cells, NRF1 is similarly regulated by GSK3-dependent phosphorylation, facilitating the degradation of NRF1 by the SCF-Fbw7 ubiquitin ligase complex (Biswas et al., 2013). NRF2 is regulated similarly by GSK3, where GSK3 phosphorylates and excludes NRF2 from the nucleus (Salazar et al., 2006) and facilitates NRF2 degradation through the SCF/β-TrCP complex (Rada et al., 2011). Activating NRF2 has been proposed as a potential therapeutic target for NGLY1 deficiency, and NRF2 can partially compensate for gene expression changes due to the dysregulation of NRF1 in NGLY1 deficient cells (Yang et al., 2018). While the interplay between NGLY1, NRF1/NRF2, and GSK3 needs further exploration, our data show that inhibiting GSK3 restores the ability of *dNGLY1* deficient flies to respond to proteasome stress.

How the modulation of the dopamine and serotonin signaling pathways rescues defects in *dNGLY1* deficient flies remains to be determined. One possibility is that the dopamine and serotonin modulators directly impact GSK3 activity. 5HT_1A_ and 5HT_1B_ receptor agonists push GSK3 towards a phosphorylated and inactive state, while agonism of the dopamine D_2_ and 5HT_2A_ receptor shifts GSK3 towards an active state (Beurel et al., 2015; Polter & Li, 2011). Therefore, GSK3 inhibition may be a key mediator of the monoamine signaling compounds identified in our screen, resulting in elevated activity of NRF1/NRF2. Indeed, we show that trimipramine rescues a proteasome inhibitor induced larval size defect in dNGLY1 deficient larvae similar to lithium and TWS119, suggesting a direct effect on NRF1/NRF2 and GSK3. The monoamine modulator aripiprazole, which has the same rescue effect as trimipramine, lithium, and TWS119, increases NRF2 levels (Iyer et al., 2019), supporting this hypothesis. Alternatively, trimipramine, and possibly other compounds like lithium, may rescue lethality and larval size defects through a separate mechanism. Trimipramine induces the expression of ER stress markers, including spliced XBP1, GRP78, CHOP, and phosphorylated IRE1a in multiple cell lines (Morita et al., 2020). Similarly, lithium treatment upregulated expression of *GRP78* and the antiapoptotic protein *Bcl-2* (Hiroi et al., 2005). Interestingly, large B-cell lymphoma cells are resistant to proteasome inhibition by bortezomib due to high levels of GRP78 expression (Mozos et al., 2011). Therefore, trimipramine and lithium may induce bortezomib resistance through the upregulation of ER stress genes, allowing flies with reduced levels of *dNGLY1* to better deal with subsequent proteasome stress. Future studies will determine whether trimipramine rescue effects that protect from proteasome stress act through GSK3 and NRF1/NRF2, ER stress, or both.

Complementary to our *in vivo* screening approach, researchers have used cell-based methods to conduct drug screens for NGLY1 deficiency therapeutics. Dactosilib, an mTOR/PI3K inhibitor, increased the degradation of NGLY1 substrates, potentially through elevated levels of autophagy (Mueller et al., 2020). In addition to regulating NRF1/NRF2, GSK3 inhibition increases autophagy (Mancinelli et al., 2017). It remains to be seen whether the GSK3 inhibitors identified in this work provide benefit in *dNGLY1* deficient flies through the mTOR pathway. An additional screen to restore morphology defects in *NGLY1^-/-^* cells identified numerous hit compounds involved in microtubule dynamics, including nocodazole (Lebedeva et al., 2021), which was also a hit in our primary screen. Although we did not focus on nocodazole in this work, future experiments will determine if nocodazole similarly rescues defects in *dNGLY* deficient flies.

In summary, we conducted an unbiased *in vivo* small molecule screen and used a gene expression-based approach to identify compounds amenable to drug repurposing for NGLY1 deficiency. This work resulted in the identification of multiple compounds that may serve as therapeutics for the NGLY1 deficiency population. Our work demonstrates the power of *Drosophila* in therapeutic development for rare diseases, and similar phenotypic, small molecule screens should be applied to other rare diseases.

## Methods

### *Drosophila melanogaster* stocks and maintenance

Stocks were maintained on standard agar-dextrose-yeast medium at 25°C on a 12-h light/dark cycle. *dNGLY1^^pl^* flies, (referred to as *dNGLY1^-^*) carry an early stop codon in the *dNGLY1* gene, and were previously characterized and generously provided by Perlara, PBC (Rodriguez et al., 2018). The *dNGLY1^-^* allele is homozygous lethal and the stock is maintained with the *CyO* balancer. *Dopa Decarboxylase-GAL4, Tyrosine Hydroxylase-GAL4, Elav-GAL4,* and *Repo-GAL4* were provided by Adrian Rothenfluh (University of Utah). The *UAS-dNGLY1^wt^* and *UAS-dNGLY1^C303A^* constructs were provided by Hamed Jafar-Nejad (Baylor College of Medicine) and described previously (Funakoshi et al., 2010). The *dNGLY1^ΔGAL4^* line was provided by Hugo Bellen (Baylor College of Medicine). The followings stocks were obtained from the Bloomington Drosophila Stock Center (Bloomington, IN): *Trh-GAL4* (BDSC # 38389), *Tubulin-GAL4* (BDSC # 5138), *UAS-dNGLY1-RNAi* (BDSC # 54853).

### Compounds

The 1040 compounds used for the small molecule screen were from the Prestwick Chemical Library (Illkirch, France). The following compounds were obtained from Cayman Chemical (Ann Arbor, MI): TWS119 (#10011251), ipsapirone (#22075), bromocriptine (#14598), urapidil (#29004), trimipramine (#15921), dichlorphenamide (#23658), nimesulide (#70640), and mesalamine (#70265). Lithium chloride (LiCl) was obtained from Sigma (St. Louis, MO, #62476), lithium acetate was obtained from Aldrich Chemical Company (Milwaukee WI, #21,319-5), and bortezomib (BTZ) was obtained from EMD Millipore (#179324-69-7).

### *In vivo* small molecule screen

Compounds from the Prestwick Chemical Library were obtained at 10 mM in DMSO and diluted to 1 mM in phosphate buffered saline. Standard Drosophila food was melted and cooled to 60°C prior to the addition of compounds at a final concentration of 5 μM. Six pre-mated *dNGLY1^+/-^ (dNGLY1^-^/CyO*) intercrossed females were placed in each vial and allowed to lay eggs for 24 hours. Visual confirmation of egg laying was confirmed prior to pre-mated female removal. The percent expected *dNGLY1^-/-^* flies was calculated based on the expected Mendelian ratios of emerging adults. The *CyO* balancer carries a wildtype *dNGLY1* allele and is homozygous lethal, so the expected ratio of *dNGLY1^+/-^* to *dNGLY1^-/-^* was 2 to 1. *dNGLY1^-/-^*flies were identified based on the absence of the CyO balancer curly wing phenotype. The *CyO* balancer also carries a GFP marker and non-curly wing flies were confirmed to lack the balancer chromosome by checking that they were also GFP negative.

### Connectivity Map Analysis

To identify potential therapeutic compounds based on gene expression profiles, we used the Connectivity Map (CMAP) online tool, available at clue.io (Lamb et al., 2006; Subramanian et al., 2017). We utilized two previously published gene expression datasets as inputs: RNAseq data from *Tubulin>dNGLY1-RNAi Drosophila* (Owings et al., 2018) and RNAseq from the liver of mice with liver-specific deletion of *Ngly1* (Fujihira et al., 2020).

### Proteasome sensitivity assay

Mated *dNGLY1^+/-^* females were placed on food containing drug or DMSO and allowed to lay eggs for approximately 8 hours. Adults were removed and after four days of development, 3^rd^ instar larva were transferred to vials containing food with DMSO, bortezomib, or bortezomib plus drug. After two days on bortezomib containing food, larvae were genotyped using the presence of GFP, as described above. Larvae were imaged at 2.5X magnification using a Leica EC3 camera. Larval size was quantified using ImageJ.

### Statistical Analysis

Statistical tests were performed using Prism version 9 (Graphpad). Correlation analysis was performed using R software with ggplot2 and ggpubr.

## Acknowledgements

This work was supported by an NIGMS R35 (R35 GM124780) and a University of Utah STARS CCTS Pilot Project award (NCATS UL1TR002538) to CYC. CYC and RTP were supported by Primary Children’s Hospital Center for Personalized Medicine. KAH was supported by an NIH/NHGRI Genomic Medicine T32 Postdoctoral Training Grant from the University of Utah (T32 HG008962) and an NIGMS F32 award (F32 GM139349). This work was also supported by a generous gift from the Might Family through the Bertrand T. Might Fellowship.

---

**Supplemental Figure S1.**
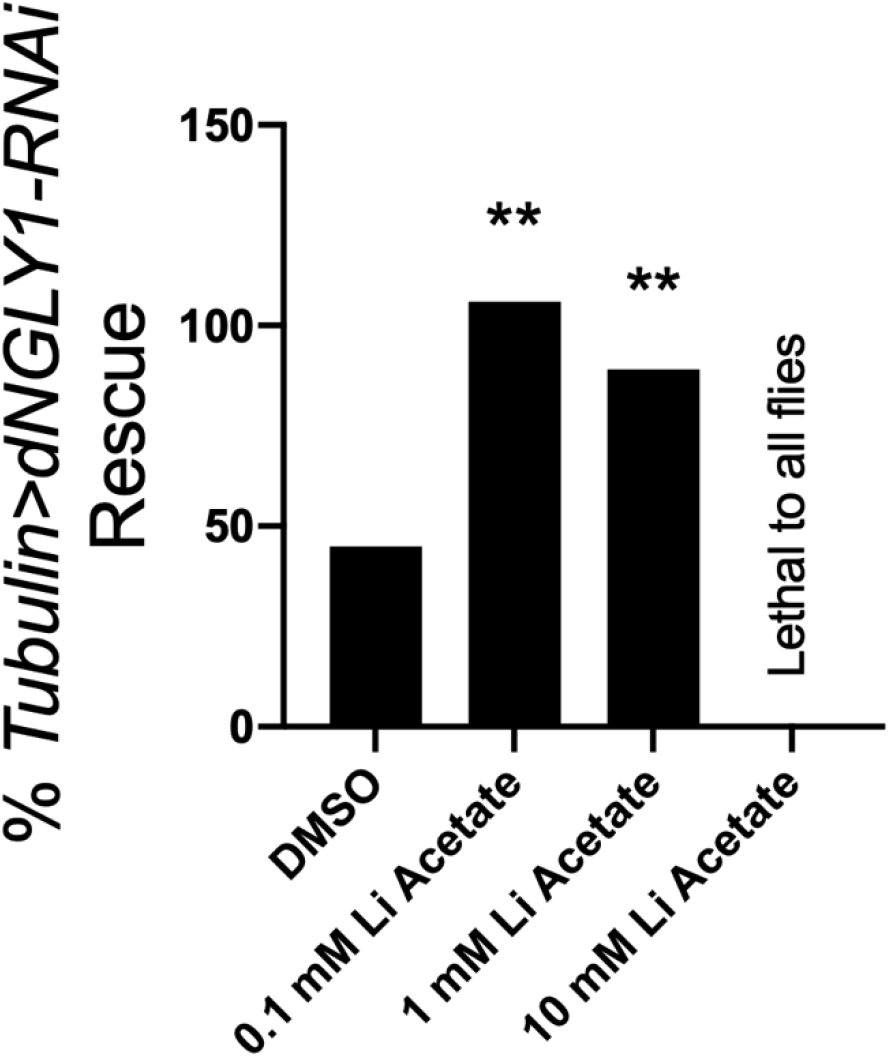
Lithium acetate rescues lethality in *Tubulin>dNGLY1-RNAi* flies. Lithium acetate rescued lethality in *Tubulin>dNGLY1-RNAi* flies at 0.1 mM and 1 mM, compared to DMSO treated controls, similar to the effects observed with lithium chloride. 10 mM lithium acetate was lethal to all flies in the cross. ** p < 0.01.

**Supplemental Figure S2.**
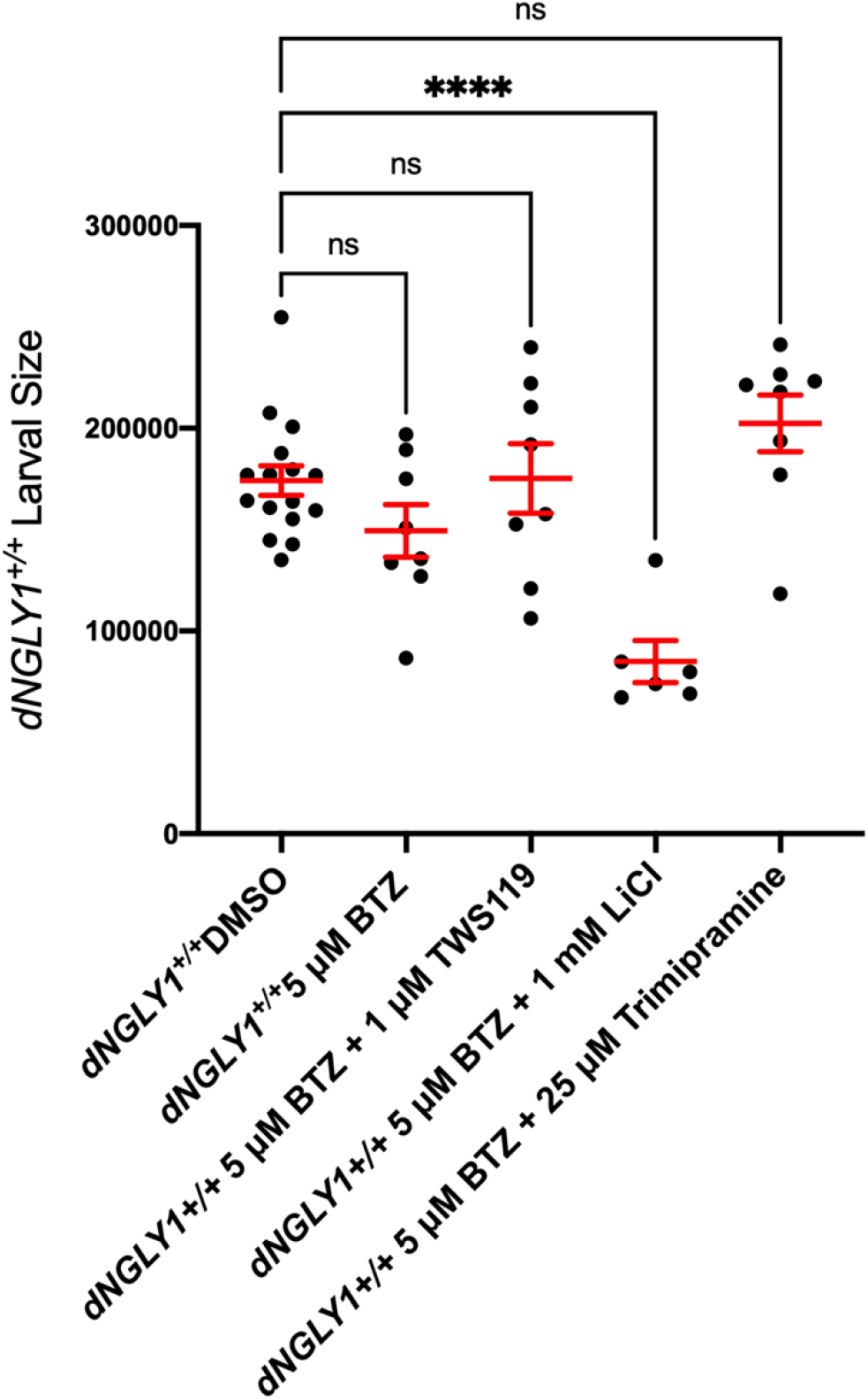
Quantification of wildtype *dNGLY1^+/+^* larva size with treatment with bortezomib (BTZ) alone or with TWS119, LiCl, or trimipramine pretreatment. There was no effect of treatment on larval size, except for BTZ + LiCl, where larvae were significantly smaller than DMSO treated controls. ****p < 0.0001.

